# *Escherichia coli*-induced gut IL-33 release inhibits lung type 2 allergic responses

**DOI:** 10.1101/2025.11.07.687219

**Authors:** Caio Loureiro Salgado, Marina Caçador Ayupe, Juliana Falcão Rodrigues, Leonardo Mandu-Gonçalves, Luisa Menezes Silva, Bárbara Cristina Pizzolante, Bernardo de Castro Oliveira, Guilherme William Da Silva, Francielly Moreira, Erick Esteves Oliveira, Marcelo Valdemir de Araújo, Gabriele Manamy Baba Rodrigues, Jofer Andree Zamame, Kristen Brammer, Igor Santiago-Carvalho, Bruna de Gois Macedo, José Carlos Alves Filho, Hirohito Kita, Elizabeth Norton, Maria Regina D’Império-Lima, Luís Carlos de Souza Ferreira, Henrique Borges Da Silva, Denise Morais da Fonseca

## Abstract

**Background:** The incidence of lung allergies is reduced in countries with higher prevalence of infection and environmental exposure to microbes. However, enteric bacterial infections do not always correlate with lower incidence of allergic disorders and how lung immunity to allergens can be regulated by gut exposure to pathogens and their toxins is not fully understood.

**Objective:** We used mouse models of enterotoxigenic *Escherichia coli* (ETEC) infection and lung allergy to examine how gut exposure to bacteria, or their related toxins, affects allergic lung inflammation.

**Methods:** Naïve C57BL/6 mice were infected with enterotoxigenic *Escherichia coli* (ETEC) or orally treated with the ETEC LT toxin, to mimic enteric bacterial infections. After two weeks, these mice were treated intranasally with Ovalbumin (OVA) and Papain or IL-33, followed by challenge with OVA, to induce allergic lung inflammation that was assessed using multiple readouts.

**Results:** Gut exposure to ETEC significantly inhibited allergic lung inflammation in a LT-dependent manner, as demonstrated by reduced tissue inflammation, less accumulation of type 2 cytokines, and reduced lung numbers of type 2 immune cells such as type 2 innate lymphoid cells (ILC2) and eosinophils. The anti-allergic capacity of LT was associated with reduced ability of lung ILC2s to recognize IL-33. Counterintuitively, deletion of either IL-33 or ILC2s significantly reverted the LT protective effect, suggesting the LT-mediated protection may occur through gut release and local sensing of IL-33.

**Conclusions:** Exposure to ETEC protects hosts against allergic lung inflammation through a negative feedback loop regulated by gut IL-33 release and sensing, suggesting a possible new immunological mechanism for reduced lung allergy incidence observed in areas with enteric bacterial infections.

**Key Messages:**

- Enteric exposure to enterotoxigenic *E. coli* (ETEC) bacteria or its toxin LT significantly protects hosts against lung type 2 allergic inflammation.
- ETEC and LT downregulate the capacity of lung ILC2s to respond to allergen-induced IL-33.
- Deletion of IL-33 or ILC2s significantly impairs the protective effect of ETEC and LT, suggesting the presence of a negative feedback loop driven by toxin-induced gut IL-33 release.

## INTRODUCTION

Allergic lung diseases, such as asthma, are driven by type 2 immune responses which in turn induce airway and parenchymal inflammation^1,2^. Together, Th2 CD4^+^ T cells and type 2 innate lymphoid cells (ILC2s) program the lung tissue for type 2 allergic inflammation, which includes the recruitment and activation of eosinophils through production of Eotaxin, IL-4, IL-5 and IL-13^3–5^. Lung-resident ILC2s play a crucial role in both the initiation and the propagation of type 2 allergic inflammation, through their ability to quickly respond to tissue-derived IL-33^6,7^. In addition to the identification of effector functions, the epidemiology of lung type 2 allergic diseases and its associated mechanisms are subjects of intense investigation. Of special interest is the inverse correlation between past microbial exposures and the development of allergic lung diseases^8,9^, which is coined the hygiene hypothesis. The incidence of lung allergies has steadily increased over the last century and the decline in microbial exposures due to post-modern lifestyle is often suggested as a causal factor ^9,10^. The definition of immune drivers of the hygiene hypothesis, however, remains incompletely understood^11–14^. Also, not all studies have reached a consensus regarding the protective relationship between enteric infection and asthma, and factors such as pathogen intrinsic features, microbiota composition, and population characteristics may further complicate this association^12–15^

While some studies on the hygiene hypothesis focused on changes in the gut microbiome^16–18^, a large body of work has indicated a fundamental role for a history of exposure to pathogens in the modulation of lung type 2 allergy^16^. Clinical studies, for example, have shown that proxies for increased pathogen exposure such as early age daycare were associated with lower allergic disease risk^19,20^. Past studies with animal models have provided additional evidence, showing that exposure to microbial products such as unmethylated CpG DNA or endotoxins decrease type 2 allergic inflammation^21–24^. Likewise, direct lung exposure to microbial products or interferon molecules, which are downstream of pathogenic exposures, decrease the ability of ILC2s to respond to IL-33^25–27^. These studies did not test, however, how physiological routes of exposure to pathogens can regulate lung type 2 allergies. A recent study used cohousing of laboratory and pet store mice to start answering this question and found that laboratory cohoused mice had a transient decrease in ILC2 responses during airway exposure to allergens^28^. This work supports the idea that recent infection exposures can suppress type 2 allergic inflammatory responses, especially ILC2s. However, there is still limited understanding about many aspects of this protective effect, especially regarding the exact type of pathogens and intrinsic factors that can drive protection.

This question is especially important considering the robust epidemiological evidence that enteric infections are inversely correlated with the incidence of allergic lung diseases in defined areas of the globe^29^. These infections are common in the tropical low-to-middle income countries and are typically caused by toxin-producing bacteria such as *Clostridium difficile* or pathogenic strains of *Escherichia coli* ^30,31^. Enterotoxigenic *E. coli* (ETEC) is one of the main causes of recurring diarrhea in infants^32^ and is a major cause of diarrhea among adults visiting endemic areas^33^. ETECs induce a gut-localized inflammatory response associated with the initial production of IL-6 and IL-1β, and the induction of gut epithelial disruption and consequent IL-33 release, followed by a tissue repair phase driven by type 2 immune responses^34^. In the case of ETECs, the heat-labile toxin (LT), especially the LT1 subunit, is a major virulence factor, participating in the disruption of gut epithelial cells^35^. Whether enteric exposure to ETECs and LT can directly modulate lung type 2 allergic inflammation is not known. This would likely involve modules of connection between the gut and lung immune responses associated with immune cell migration between these organs^36,37^, downstream effects of the local release of cytokines^38^, or in some cases the systemic release of soluble factors^39,40^. Defining the mechanisms by which enteric exposure to bacterial toxins could alter immune responses outside of this tissue is, therefore, another unknown subject.

Here, we describe that gut exposure to ETEC in or LT-dependent manner constitutes a non-classical pathogen-mediated mechanism for controlling allergic lung inflammation. Specifically, we used mouse models of oral gavage and found that mice exposed to ETEC or LT were significantly protected from lung type 2 allergic inflammation. In contrast with a gut-local increase in IL-33 release, oral exposure to ETEC or LT led to a reduced ability of lung ILC2s to respond to IL-33, evidenced by both their downregulation of the IL-33 receptor ST2 and the lack of response to intranasal IL-33 challenge. Strikingly, this protective effect is significantly reverted in mice lacking IL-33 and ILC2s. Altogether, these results suggest that exposure to enteric bacteria and the local release of IL-33 induces a distal negative feedback effect on lung type 2 immune response capacity, promoting protection against lung type 2 allergy. Our data provide a mechanistic basis for how enteric bacterial infections can inhibit the development of allergic lung diseases and may help explain the inverse correlation in the incidence of diarrhea-inducing defined infections and asthma.

## METHODS

### Mice

Eight-week-old female C57BL/6 (WT-Wild type), IL-10 and IFN-γ knockout (KO) were obtained from the Immunology Department (Institute of Biomedical Sciences, University of Sao Paulo) animal facility or from Jackson Laboratories. IL-33 green-fluorescent protein (GFP) knock-in (IL-33KO) mice were provided by José Carlos Alves Filho (University of Sao Paulo). Villin-Cre and *Il33*^fl/fl^ mice were obtained from Jackson Laboratories, then cross-bred in house. *Il7r*-Cre x *Rora*^fl/fl^ mice (ILC2KO) were provided by Hirohito Kita (Mayo Clinic Arizona). Mice were maintained under specific pathogen-free conditions, with alternating 12 hours of light/dark cycles and fed a regular diet and water *ad libitum*. Experiments were conducted in accordance with the Ethics Committee of the University of São Paulo (CEUA-USP, 8229211019) and IACUC-Mayo Clinic (protocols A00005172-20-R23, A00005542-20-R23).

### ETEC infection

Enterotoxigenic *E. coli* (Strain PE0262 O8:H9)^41^, producing LT1 type only and isolated from diarrheic subject, and one plasmid-cured non-toxigenic derivative (LT-KO) of the ETEC 4611-4 (O159:H21)^41^ strains were included in the present study. The ETEC strains were grown in Tryptic soy broth (TSB) media (Kasvi) for 3 h, at 37°C, 200 rpm. C57BL/6 mice were treated with streptomycin (5 g/L, diluted in water)^42^ via gavage two days before infection with 10^9^ colony-forming units (CFU) of LT-producing or LT-KO ETEC via oral gavage.

### Quantification of ETEC infection

During the infection period, fecal samples were collected from the animals, weighed, and serially diluted in sterile phosphate-buffered saline (PBS). The dilutions were plated onto Tryptic soy agar (TSA - Kasvi) supplemented with streptomycin (Sigma-Aldrich) (50 µg/mL) to select for the infecting *E. coli* strain. Plates were incubated overnight at 37°C, and colonies were enumerated to determine colony-forming units (CFU) per gram of feces. For tissue translocation assays, animals were euthanized at selected time points. The spleen, liver, and lungs were aseptically collected; each tissue was weighed and homogenized in sterile PBS using. Homogenates were serially diluted and plated onto TSA containing streptomycin (50 µg/mL), followed by incubation overnight at 37 °C. Stool and tissue DNA were analyzed for the ETEC-specific LT gene to determine the level of organism shedding in stool and the bacterial burden in the tissues^43^.

### Protein purification

LT, LTB or LTK63 proteins were purified through affinity chromatography using Galactose-gel resin (Pierce, Immobilized D-Galactose Gel) packed in an XK 16/20 column from Amersham Biosciences. The automated purification process was carried out using an AKTA - Pure apparatus (GM). Column-based purification of LT proteins was done as described before^44^. Quantification of purified protein concentration was performed by measuring absorbance on a spectrophotometer (Gene Quant spectrophotometer GE Amersham Biosciences), following a previously described protocol^45^.

### LT exposure

C57BL/6 mice were orally treated with 2.5% sodium bicarbonate to neutralize stomach pH. After 15 min, LT toxin administration was performed via oral gavage. The animals were immunized with 0.6 mg/kg doses for the first 3 days, followed by 0.3 mg/kg µg doses for 5 days with a 1-day interval between treatments.

### Allergy induction

Mice were intranasally immunized 3 times with Ovalbumin (Sigma-Aldrich) (OVA, 50 μg) and papain (Sigma-Aldrich) (Pap, 20 μg) (OVA+Pap) with 2-day intervals. Seven days after the last immunization, 2 challenges were carried out with OVA (50 μg) intranasally, with 7 days between each challenge^46^. Mice were euthanized 48 h after the second challenge. In other experiments, mice were treated intranasally with recombinant IL-33 (R&D Systems) 0.1 µg/per animal for 3 days, with a 1-day interval^47.^

### Tissue isolation and processing

Prior to tissue collection, anti-CD45 was injected intravenously to discriminate between vascular-associated and tissue-associated leukocytes, as described previously^48^. Small intestines were collected and maintained in RPMI (Roswell Park Memorial Institute Medium, Sigma-Aldrich, St Louis MO) (Sigma-Aldrich) supplemented with 3% of FBS (Fetal Bovine Serum – LGC Biotechnology). The mucus was removed with RPMI and EDTA (2 mM; LGC Biotechnology) solution, through vigorous shaking. After this process, to disrupt the intraepithelial junctions, a solution containing RPMI, EDTA (5 mM), 3% FBS, DTT (Dithiothreitol, 0.145 mg/mL; Sigma-Aldrich) was added to tissue, followed by incubation at 25 min at 37°C and 180 rpm or 4 g. After the incubation, the suspension was filtered and RPMI containing 2 mM EDTA was added; this was repeated three times. Fragments were cut into small pieces and resuspended with RPMI medium containing collagenase IV (0.75 mg/mL; Sigma-Aldrich) and DNase I (0.5 mg/mL; Sigma-Aldrich) solution, followed by incubation at 25 min, at 37°C and 4 g. The content was filtered in a 100 µm cell strainer, centrifuged (10 min, at 4°C and 1600 rpm or 315 g), and filtered in 70 µm and 40 µm cell strainers. Cells were resuspended in complete RPMI medium containing 2% Penicillin/streptomycin (HyClone), 1% Pyruvate (Sigma-Aldrich), 0.25% β-mercaptoethanol (SigmaAldrich), 2% L-Glutamine (SigmaAldrich), 5% FBS (Sigma-Aldrich) and 1% non-essential amino acids (HyClone). Cell viability was counted in a Neubauer chamber in the presence of trypan blue dye and cells were stained for flow cytometry.

For lung cell isolation, lungs were sliced into small fragments and enzymatically digested using collagenase IV (1 mg/mL) (Sigma-Aldrich) and DNase I (0.5 mg/mL), for 30 min, incubation at 37°C with agitation at 180 rpm or 4 g. Cells were suspended in RPMI supplemented with 3% FBS, filtered through a 70 µm cell strainer, and centrifuged (10 min/4°C/1600 rpm or 315 g). Cell pellets were then resuspended in complete RPMI medium. Samples were maintained on ice, and cell viability was assessed using trypan blue dye in a Neubauer chamber. Samples were then stained for flow cytometry.

### Surface antibody staining for flow cytometry

Cell suspensions (1-2 × 10^6^ cells/sample) were transferred to U-bottom 96-well plates, washed with Hanks’ Balanced Salt Solution (HBSS; Gibco) by centrifugation (3 min, at 4°C and 1600 rpm or 315 g). Cell viability dye Live/Dead Cell Stains (LD, Thermo Fisher Scientific) was used for 20 min at 4°C, protected from light. Subsequently, cells were washed with FACS buffer (PBS / 2% FBS) and resuspended in 30 µL of the surface antibody mix **(Supplementary Table 1)**, together with Fc-block (anti-CD16/32) and incubated for 30 min at 4°C, protected from light. After this period, cells were washed with FACS buffer and fixed in 2% paraformaldehyde (Sigma-Aldrich), 20 min at room temperature – RT and resuspended in FACS Buffer. Samples were acquired on the flow cytometer LSR FortessaX-20 (BD Biosciences) or Aurora C5 (Cytek), and all analyses were performed with FlowJo® v10 software (BD Biosciences).

### Intracellular antibody staining for flow cytometry

Cell suspensions (1-2×10^6^ cells/sample) were aliquoted into 96-well plates. To evaluate cytokine production, the cells were stimulated with phorbol myristate acetate (PMA) (100 µg/mL) (Sigma-Aldrich), ionomycin (1 mg/mL) (Sigma-Aldrich), and brefeldin A/Golgi Plug (BD Biosciences) in complete RPMI for 4 h at 37°C in a CO_2_ chamber. Subsequently, the cells were washed with HBSS and stained with LD and surface antibodies. Cells were then washed with FACS buffer and resuspended in fixation/permeabilization buffer - Cytofix/Cytoperm (BD Biosciences) for 20 min at 4°C protected from the light. Following washing with PermWash (BD Biosciences), cells were resuspended in the intracellular antibody mix (IL-5 and IFN-γ) and incubated for 1 h at 4°C, protected from light. Subsequently, samples were analyzed using LSR Fortessa X-20 (BD Biosciences) or Aurora C5 (Cytek) cytometers, and all analyses were conducted using FlowJo® software (BD Biosciences).

### ELISA cytokine measurement

Lung cytokine quantification was determined using multiplex ELISA (Millipore, MCYTOMAG-70K) following the manufacturer’s protocol.

### Histological analyses

The left lobes of mice lungs were harvested and fixed in 10% formaldehyde (Synth, Brazil). The specimens were then paraffin-embedded, sectioned at 5 µm, followed by hematoxylin and eosin (HE) or periodic acid-Schiff (PAS) staining. For the determination of the inflammatory score, HE-stained slides were examined in their entirety using a Leica optical microscope at 200X magnification. The peribronchiolar and perivascular areas were assessed based on the degree of inflammatory cellular infiltrate, with scores ranging from 0 to 5 for each observed field: 0) Absence of inflammatory process; 1) Sparse inflammatory cells; 2) Monolayer of inflammatory cells around the evaluated structure; 3) Ring of inflammatory cells containing 2 to 3 layers; 4) Focal cluster of inflammatory cells around the structure; 5) Intense inflammatory infiltrate. The mucosal area was measured in 3 random fields of each lobe using ImageJ software.

### Lung respiratory capacity analyses

After the last OVA+Pap intranasal challenge in the asthma model (48 h), mice were anesthetized with Pentobarbital (Nembutal; 50 mg/mL) and placed in the FlexiVent system (SCIREQ Scientific Respiratory Equipment Inc.), where respiratory resistance was measured by instilling mice with methacholine (Sigma-Aldrich) at a dose of 100 mg/mL. Mice were ventilated at a rate of 90 breaths/min with a tidal volume of 20 mL/kg during nebulization and otherwise at 150 breaths/min with a tidal volume of 20 mL/kg while breathing on the equipment. After measurements, mice were euthanized with an overdose of Pentobarbital.

### Quantification and statistical analyses

Details on statistics used can be found in figure legends. Statistical differences were calculated by using unpaired two-tailed Student’s t-test, one-way ANOVA with Tukey post-test, two-way ANOVA with Bonferroni’s post-test, or Kaplan-Meier survival curve analysis. All experiments were analyzed using Prism 9.2.0 (GraphPad Software). Graphical data was shown as mean values with error bars indicating the SD. P values of < 0.05 (*), < 0.01 (*), < 0.001 (***) or < 0.0001 (****) indicated significant differences between groups.

## RESULTS

### Gastrointestinal infection by ETEC promotes the accumulation of type 2 immune cells in the lung

First, to understand the impact of ETEC infection in the gut-lung axis, we conducted a kinetic study of infection with *E. coli* enterotoxigenic strain PE0262 in WT mice (**Figure 1A**), analyzing different immune cell populations – gating strategies are found in **Supplementary Figure 1**. On days 3, 15, and 35 p.i. (post-infection), no changes were observed for total leukocyte numbers in the small intestine lamina propria (siLP; **Supplementary Figure 2A**). However, the frequency and total number of siLP eosinophils significantly increased 35 days after infection (**Figures 1B-C**). We also observed an increase in siLP Th2 CD4+ T cells during the early stages of infection (**Figure 1D**), though without significant differences in IL-5-producing Th2 cells (**Figure 1E**). Finally, siLP ILC2s accumulated and are sustained after the clearance of infection (**Figure 1F**), accompanied by elevated percentages of IL-5+ ILC2s (**Figure 1G**) and expression of the activation marker CD44^49^ (**Figure 1H**). Therefore, we conclude that gastrointestinal ETEC infection promoted type 2 immune responses in the small intestine, which is sustained after the clearance of bacteria.

**Figure 1.**
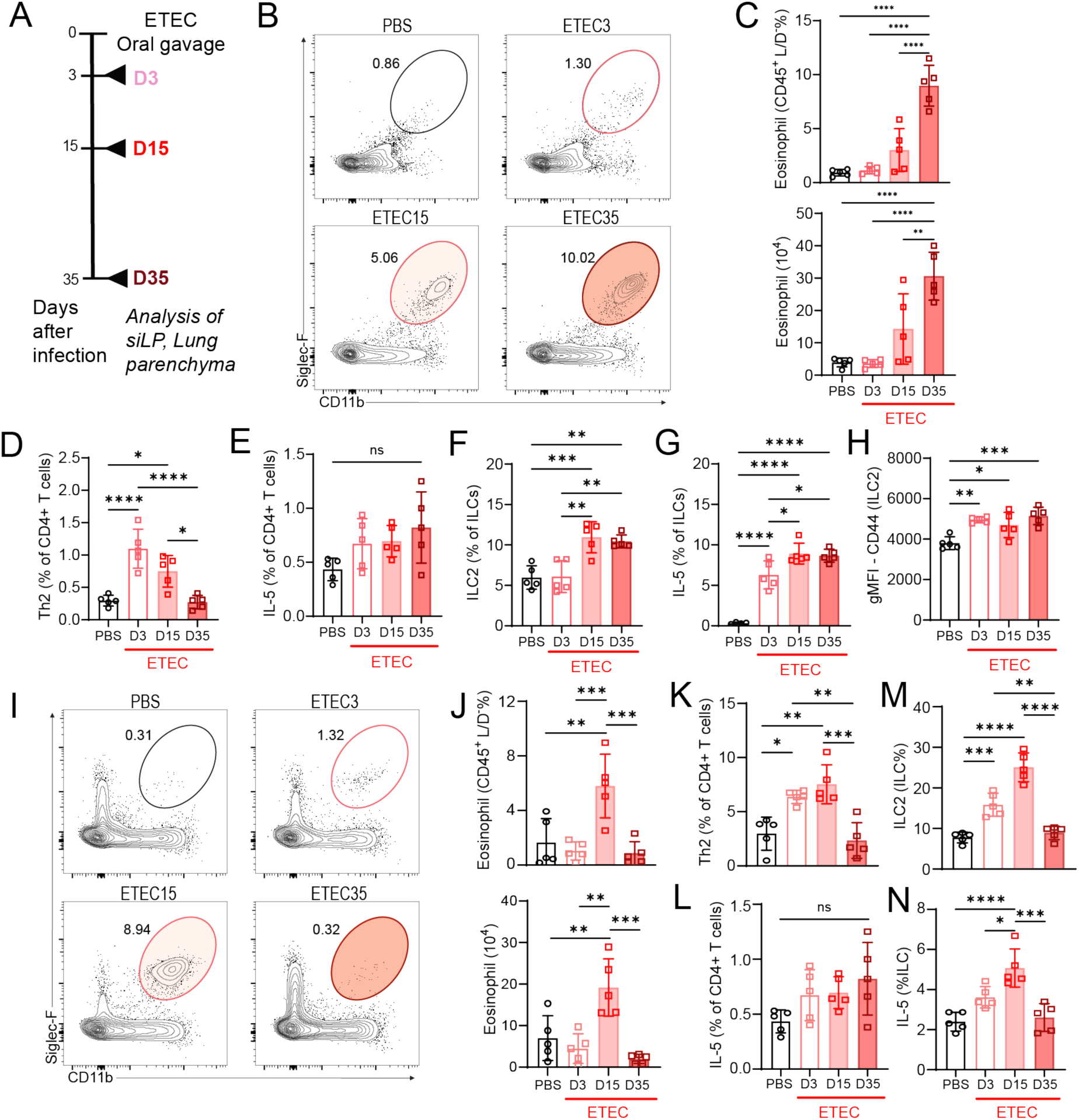
ETEC gastrointestinal infection induces the accumulation of type 2 immune cells in the small intestine and lung. (A) Representative scheme of the kinetics of infection and subsequent analyses of mice infected with LT-producing ETEC strain. (B) Representative flow cytometry plots showing small intestine lamina propria (siLP) eosinophils. (C) Average percentages and numbers of siLP eosinophils. (D) Average percentages of siLP CD4+ Th2 cells. (E) Average percentages of siLP IL-5+ CD4+ T cells. (F) Average percentages of siLP ILC2s. (G) Average percentages of siLP IL-5+ ILC2s. (H) Average geometric mean fluorescent (gMFI) values for CD44 in siLP ILC2s. (I) Representative flow cytometry plots showing lung eosinophils. Percentages referent to the eosinophil gate shown in Supplementary Figure 1 are shown. (J) Average percentages and numbers of lung eosinophils. (K) Average percentages of lung parenchyma CD4+ Th2 cells. (L) Average percentages of lung parenchyma IL-5+ CD4+ T cells. (M) Average percentages of lung ILC2s. (N) Average percentages of lung parenchyma IL-5+ ILC2s. Data are shown as means ± SD (n = 4 or 5 mice) from 2 independent experiments. One-way ANOVA with Tukey’s multiple comparison test. ****P < 0.0001, ***P < 0.001, **P < 0.01, *P < 0.05.

ETEC remains localized to the intestines in mice, during one week after infection, and did not translocate to distal tissues such as the lung (**Supplementary Figure 2B**). No changes in the total number of leukocytes in the lung parenchyma were observed (**Supplementary Figure 2C**). However, a transient increase in lung eosinophil frequencies and numbers following gut ETEC infection was found (**Figure 1I-J**). Lung eosinophilia was accompanied by a significant increase in Th2 cells (**Figure 1K**), although no differences were detected in IL-5 production by CD4+ T cells (**Figure 1L**). A similar increase was observed for lung ILC2s (**Figure 1M**), along with their ability to produce IL-5 and to express higher levels of CD44 (**Figure 1N, Supplementary Figure 2D**). Thus, gastrointestinal ETEC infection leads to an increase in lung-resident type 2 immune cells. Of note, this response was transient and was reversed at 35 days post-infection.

### ETEC infection protects from allergic lung inflammation

Because ETEC infection led to an increase in lung type 2 immune cells, we postulated that ETEC-exposed mice would have increased susceptibility to type 2 lung allergy. To test that hypothesis, we sensitized ETEC-exposed mice to intranasal OVA+Pap (**Figure 2A**). Surprisingly, ETEC infection promoted increased resistance to OVA+Pap-induced allergic inflammation. In the bronchoalveolar lavage (BAL) of ETEC-infected mice there was reduced inflammatory total cell and eosinophilic infiltration (**Figures 2B-C**), which was accompanied by a decrease in cellular layers and scattered inflammatory cells, as well as a reduction in mucus-filled areas within the alveolar lumen (**Figures 2C-E**).

**Figure 2.**
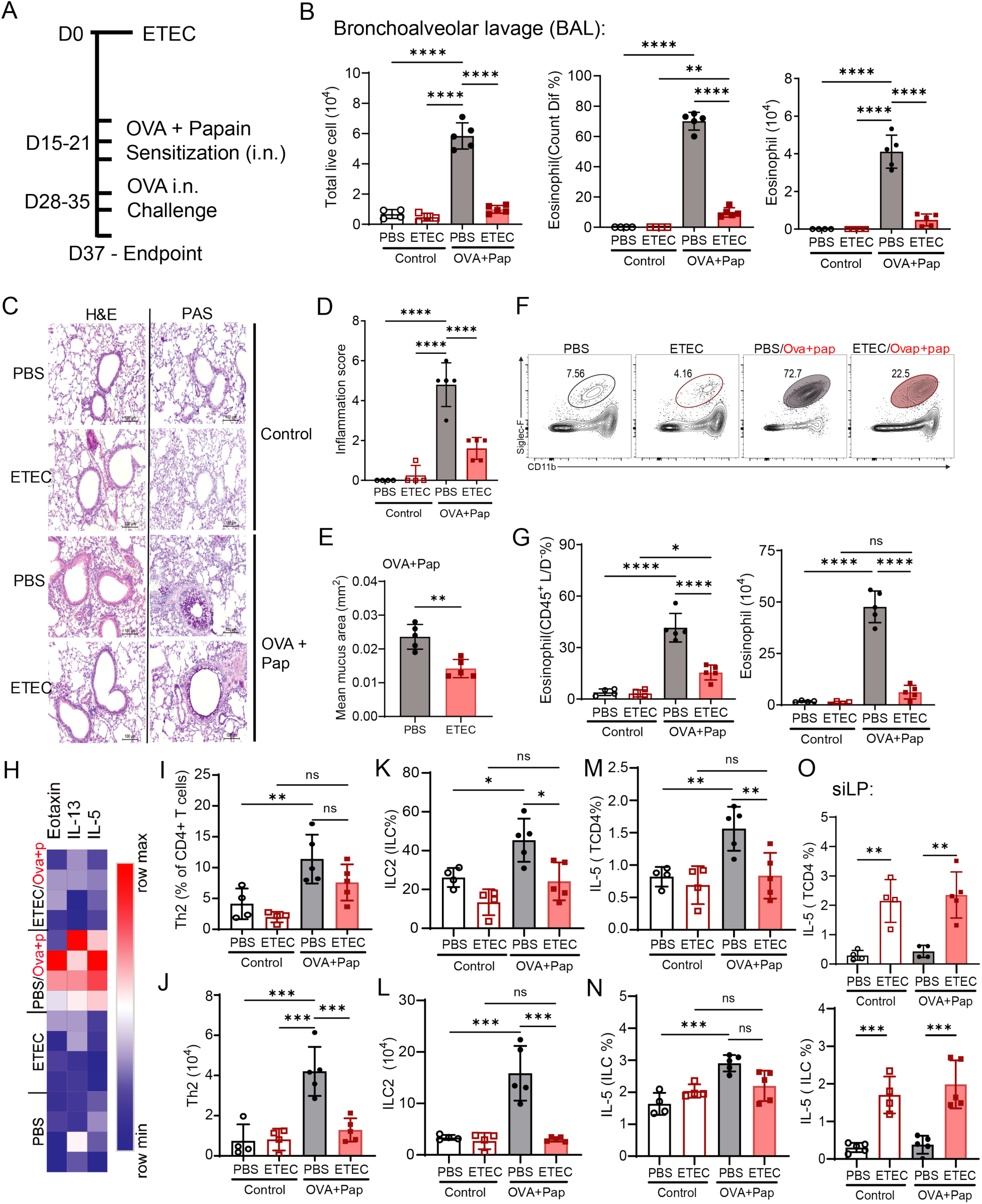
Gastrointestinal ETEC infection attenuates the development of lung type 2 allergic inflammation. (A) Representative scheme of the induction of lung allergy with OVA+Pap after ETEC infection. (B) Average numbers of live leukocytes (CD45+ L/D-) and average frequencies and numbers of eosinophils in the lung bronchoalveolar lavage (BAL). (C) Representative lung sections stained with HE and PAS. (D) Average lung inflammation score values based on histological analyses. (E) Average mean lung mucus area values based on histological analyses. (F) Representative flow cytometry plots showing control and OVA+Pap lung eosinophils. Percentages referent to the eosinophil gate shown in Supplementary Figure 1 are shown. (G) Average percentages and numbers of lung parenchyma (CD45 i.v.-) eosinophils. (H) Heatmap of cytokine levels (Eotaxin, IL-13, IL-5) measured in lung tissues using multiplex ELISA (Millipore). (I) Average percentages of lung parenchyma CD4+ Th2 cells. (J) Average numbers of lung parenchyma CD4+ Th2 cells. (K) Average percentages of lung parenchyma ILC2s. (L) Average numbers of lung parenchyma ILC2s. (M) Average percentages of lung parenchyma IL-5+ CD4+ Th2 cells. (N) Average percentages of lung parenchyma IL-5+ ILC2s. (O) Average percentages of siLP IL-5+ CD4+ Th2 cells and ILC2s. Data are shown as means ± SD (n = 4-5 mice) from 3 independent experiments. One-way ANOVA with Tukey’s multiple comparison test. ****P < 0.0001, ***P < 0.001, **P < 0.01, *P < 0.05.

In line with the BAL findings, previously ETEC-infected mice showed reduced lung accumulation of leukocytes (**Supplementary Figure 3A**) and eosinophils (**Figures 2F-G**). Moreover, pre-exposure to ETEC led to a reduced production of Eotaxin, IL-13, and IL-5 in response to OVA+Pap (**Figure 2H**). Likewise, Th2 CD4+ T cell (**Figures 2I-J, M**) and ILC2 (**Figures 2K-L, N**) responses were reduced in previously ETEC-infected mice. This reduction of type 2 responses was lung-localized, since ETEC-exposed mice maintained their type 2 gut immune signatures in the presence or absence of intranasal allergic challenge (**Figures 2O, Supplementary Figure 3B-C**). ETEC-induced protection was unlikely to be due to an induction of type 1 immune responses or anti-inflammatory IL-10 production, since bacteria-exposed IFNγKO and IL-10KO mice were still protected from OVA+Pap allergy induction (**Supplementary Figure 4**). Thus, gut-localized ETEC infection impedes the development of lung type 2 inflammatory responses induced by allergens.

### LT deletion abolishes ETEC-mediated protection from allergic lung inflammation

ETEC bacteria express multiple virulence factors to support their biological function, and the LT toxin is a virulence factor reportedly involved in cell invasion^50^. To test if LT expression (and therefore the capacity for cell invasion) is necessary for the ETEC-mediated protection from lung allergy, we infected mice with LT-KO ETEC and subsequently exposed these mice to OVA+Pap (**Figure 3A**). In contrast to WT-ETEC, LT-KO ETEC pre-exposure did not lead to reduced type 2 inflammation following OVA+Pap. That was evidenced by high total cellular and eosinophilic infiltration in the BAL (**Figure 3B**), as well as high accumulation of total lung leukocytes (**Figure 3C**). Eosinophil numbers in the lung tissue were also higher in these mice, compared to the respective non-allergic controls (**Figures 3D-E**), as well as Th2 and ILC2 (**Figure 3F-G**). These data indicate that LT expression by ETEC is necessary for their ability to inhibit type 2 allergic lung inflammation.

**Figure 3.**
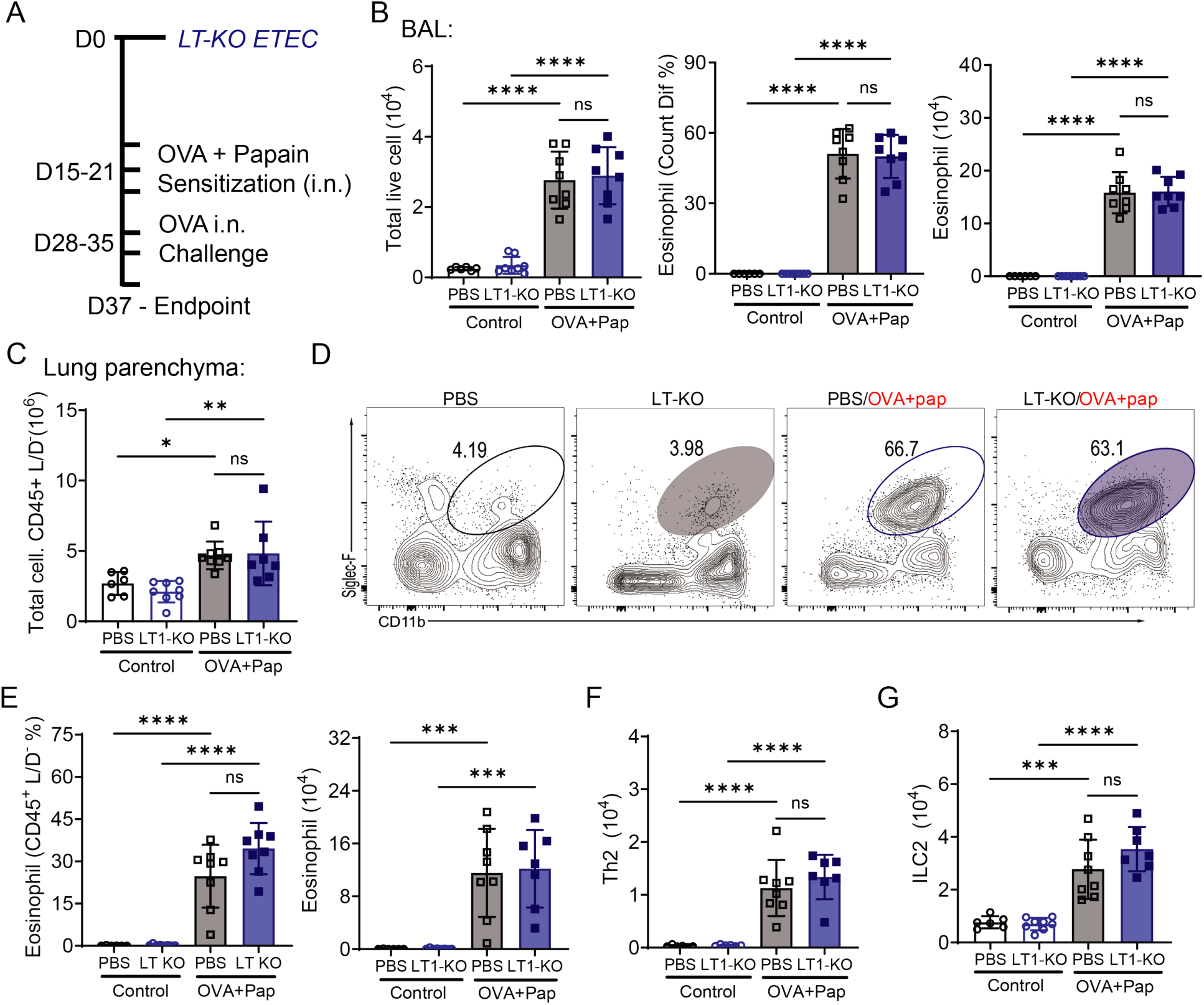
LT-deficient ETEC does not protect from allergic lung inflammation. (A) Representative scheme of the induction of lung allergy with OVA+Pap after LT-KO ETEC infection. (B) Average numbers of total leukocytes, frequency and numbers of eosinophils in the BAL. (C) Average numbers of lung total leukocytes. (D) Representative flow cytometry plots showing control and OVA+Pap lung parenchyma eosinophils. Percentages referent to the eosinophil gate shown in Supplementary Figure 1 are shown. (E) Average frequency and numbers of lung parenchyma eosinophils. (F) Average numbers of lung parenchyma CD4+ Th2 cells. (G) Average numbers of lung parenchyma ILC2s. Data are shown as means ± SD (n = 6-9 mice) from 2 independent experiments. One-way ANOVA with Tukey’s multiple comparison test. ****P < 0.0001, ***P < 0.001, **P < 0.01, *P < 0.05.

### Oral administration of purified LT is sufficient to impede lung type 2 allergic inflammation

Since the ETEC protective effect against lung allergy was dependent on LT, we tested if ETEC-derived proteins could promote the same anti-allergic effects as ETEC infection. Different recombinant ETEC toxins were produced, and mice were orally exposed to them. LTB is involved in the process of LT binding to epithelial cells through the GM1 ganglioside (monosialotetrahexosylganglioside) and LTK63 is the non-toxic variant of the LT toxin, both recombinant proteins being used as vaccine adjuvants to induce a pro-inflammatory response^51^. We observed that LT, but LTB and LTk63, induced gut type 2 immune responses similar to ETEC infection (**Supplementary Figure 5**). We then treated LT-exposed mice with intranasal OVA+Pap (**Figure 4A**). In line with ETEC findings, mice pre-exposed to LT protein had significantly reduced allergic lung inflammatory responses. This included decreased BAL accumulation of total cells and eosinophils (**Figure 4B**), as well as decreased inflammation and mucus accumulation in the lung parenchyma (**Figures 4C-E**). In addition, LT-exposed mice had decreased inflammatory accumulation of lung parenchyma total leukocytes, eosinophils, CD4+ Th2 cells and ILC2s, together with reduced IL-5 production by type 2 lymphocytes (**Figures 4F-K, Supplementary Figure 6A**). Also, similarly to ETEC infection, the LT-mediated protection was lung-localized since type 2 immune responses in the gut, particularly those ILC2-mediated, were still present after intranasal OVA+Pap (**Supplementary Figure 6B**). These data added evidence that gut exposure to ETEC-derived LT protein inhibits type 2 immune responses to lung allergens.

**Figure 4.**
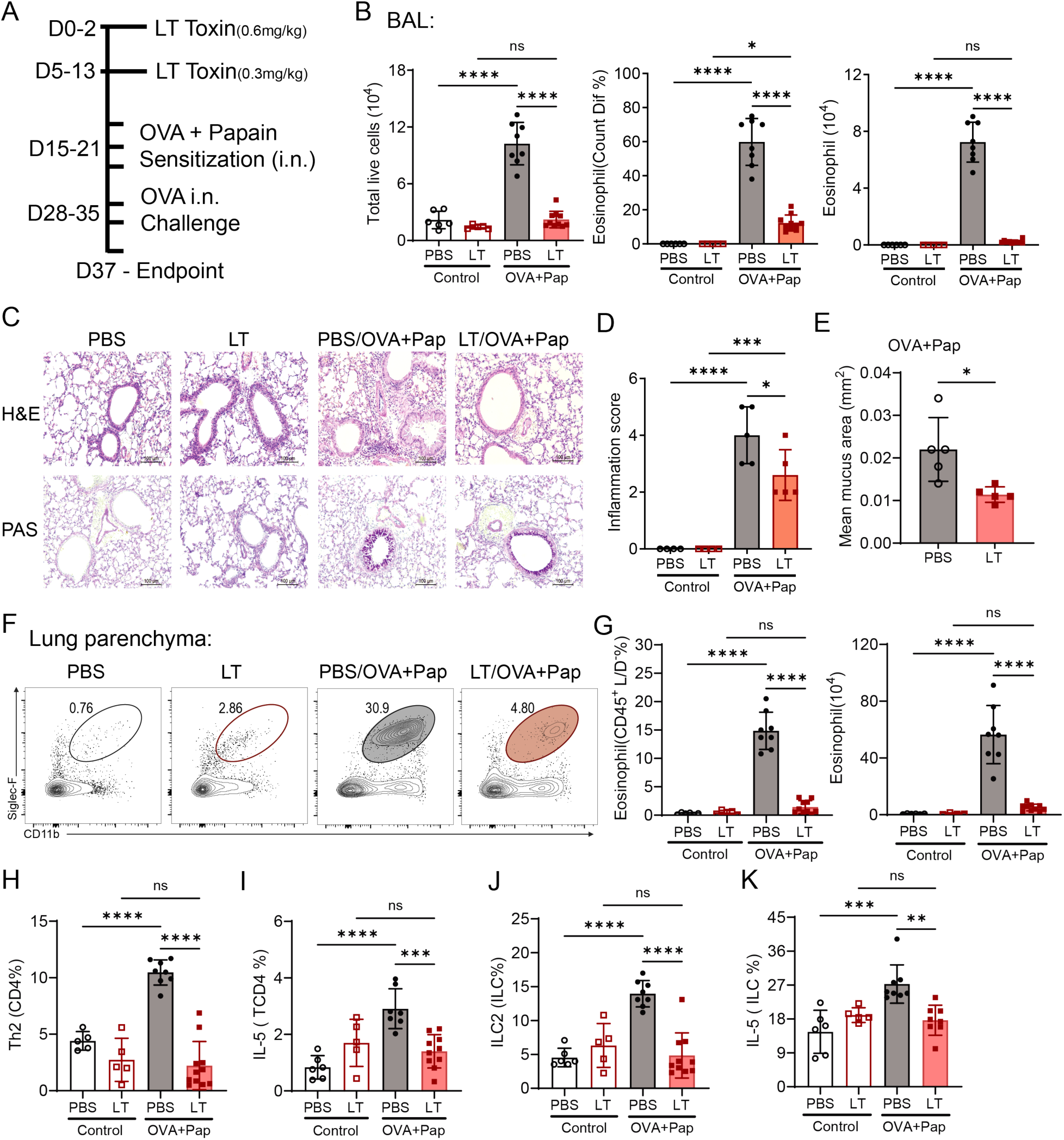
Gut exposure to LT toxin is sufficient to promote protection from allergic lung inflammation. (A) Representative scheme of the induction of lung allergy with OVA+Pap after oral gavage with recombinant LT protein. (B) Average numbers of total leukocytes, frequency and numbers of eosinophils in the BAL. (C) Representative lung sections stained with HE and PAS. (D) Average lung inflammation score values based on histological analyses. (E) Average mean lung mucus area values based on histological analyses. (F) Representative flow cytometry plots showing control and OVA+Pap lung parenchyma eosinophils. Percentages referent to the eosinophil gate shown in Supplementary Figure 1 are shown. (G) Average frequency and numbers of lung parenchyma eosinophils. (H) Average numbers of lung parenchyma CD4+ Th2 cells. (I) Average frequency of IL-5+ lung parenchyma CD4+ T cells. (J) Average frequency of lung ILC2s. (K) Average frequency of lung parenchyma IL-5+ ILC2s. Data are shown as means ± SD (n = 5 - 9 mice) from 3 independent experiments. One-way ANOVA with Tukey’s multiple comparison test. ****P < 0.0001, ***P < 0.001, **P < 0.01, *P < 0.05.

### Gastrointestinal LT exposure decreases the ability of lung immune cells to respond to IL-33

Next, we examined the immune events occurring in the gut following ETEC/LT exposure. The LT-mediated invasion of gut epithelial cells by ETEC can induce cell death^52^; a common by-product of epithelial cell death is the release of inflammatory IL-33, which is a danger signal and amplifies local immune type 2 responses^53^. Indeed, there was an increase in IL-33-producing gut epithelial cells following ETEC infection, while no differences were observed for lung epithelial cells (**Figure 5A**). Associated with this response, we observed a sustained increase in the expression of the IL-33 receptor ST2 in gut ILC2s from mice exposed to oral LT (**Figure 5B**). Paradoxically, lung ILC2s – despite increased accumulation in response to LT (**Supplementary Figure 6C**) – display a significant reduction in ST2 expression (**Figure 5C**).

**Figure 5:**
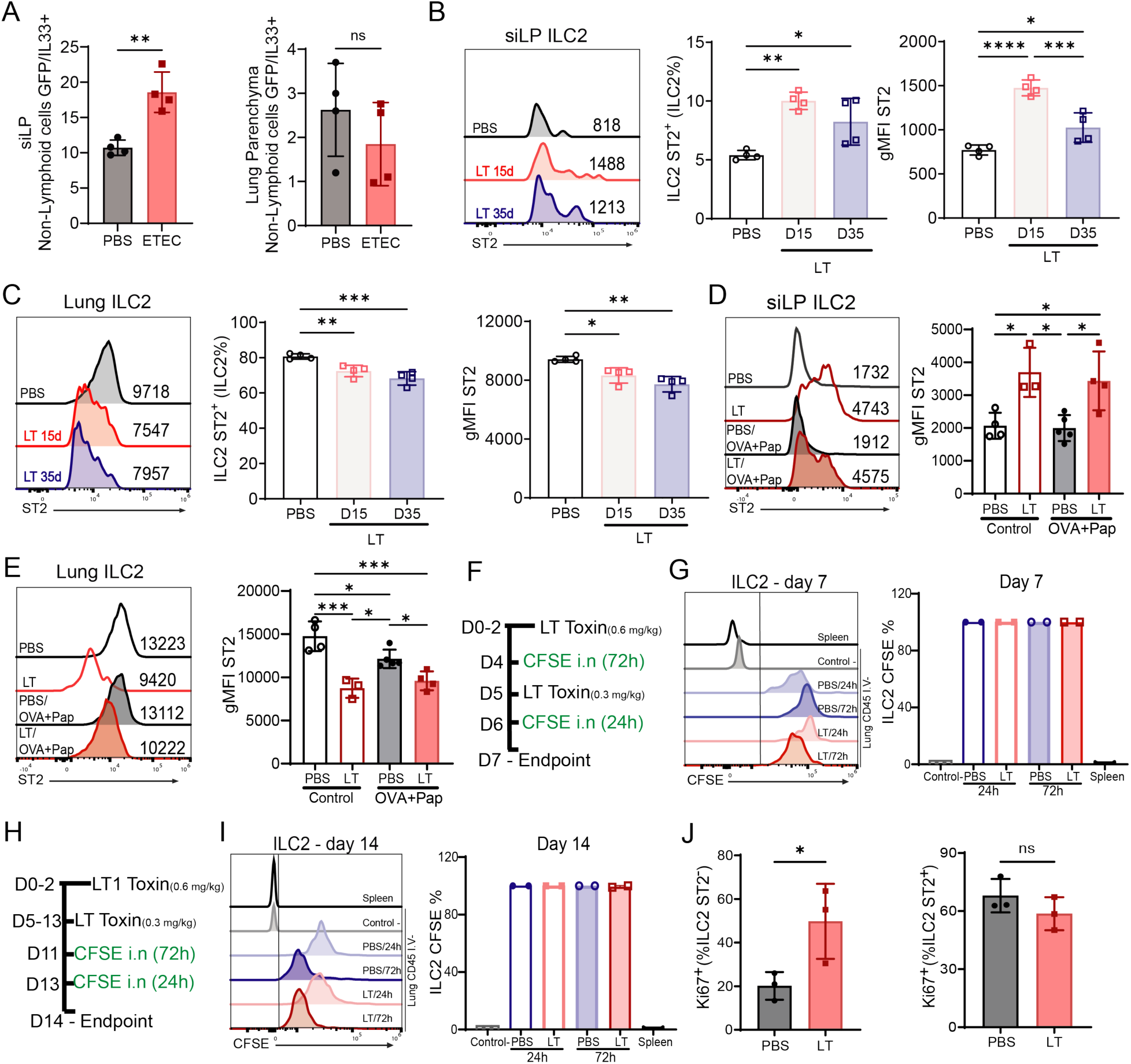
Gastrointestinal LT exposure leads to decrease in ST2 expression by lung ILC2s. (A) Average frequency of non-lymphoid cells (CD45-) positive for IL-33 (IL-33-GFP+) in the gut and lung parenchyma. (B) Quantification of ST2 expression by siLP ILC2s. In the left, representative flow cytometry plots showing ST2 expression by ILC2s; in the middle, average frequency of ST2+ ILC2s; in the right, average gMFI values for ST2 in siLP ILC2s. (C) Quantification of ST2 expression by lung ILC2s. In the left, representative flow cytometry plots showing ST2 expression by ILC2s; in the middle, average frequency of ST2+ ILC2s; in the right, average gMFI values for ST2 in lung parenchyma ILC2s. (D) Representative flow cytometry plots (left) and average gMFI values (right) of ST2 in control versus OVA+Pap siLP ILC2s, exposed or not with gut LT. (E) Representative flow cytometry plots (left) and average gMFI values (right) of ST2 in control versus OVA+Pap lung ILC2s, exposed or not with gut LT. (F) Representative scheme of the experiments involving intranasal CFSE for short-term fate mapping of lung immune cells at 7 days of LT-exposure. (G) Representative flow cytometry plot showing CFSE staining by ILC2s at day 7 of LT-exposure (left) and Average frequency of CFSE+ lung parenchyma ILC2s (right). (H) Representative scheme of the experiments involving intranasal CFSE for short-term fate mapping of lung immune cells at 14 days of LT-exposure. (I) Representative flow cytometry plot showing CFSE staining by ILC2s on day 14 at LT-exposure (left) and Average frequency of CFSE+ lung parenchyma ILC2s (right). (J) Quantification of Ki67 expression by lung parenchyma ILC2s ST2- and ST2+ at 21 days of LT-exposure. Data are shown as means ± SD (n = 2 - 9 mice) from 2 independent experiments. One-way ANOVA with Tukey’s multiple comparison test. ****P < 0.0001, ***P < 0.001, **P < 0.01, *P < 0.05.

We next evaluated the expression pattern of ST2 by ILC2s from mice pre-exposed to LT and sensitized with intranasal OVA+Pap. Gut ILC2s retained high expression of ST2 (**Figure 5D**), regardless of intranasal allergen exposure. In the opposite direction, lung ILC2 average ST2 expression remained lower in LT-exposed mice, even after OVA+Pap exposure (**Figure 5E**). We reasoned that this lower average ST2 expression in lung ILC2s could be due to the influx of ST2-ILC2s from other sites^54,55^. To test that, we used intranasal CFSE labeling to perform a short-term fate mapping of lung immune cells^55^ (**Figure 5F - H**). Contrary to our original postulation, lung ILC2s from LT-exposed (7 and 14 days) mice did not originate from other organs (**Figure 5G - I**). In LT-exposed mice, lung ILC2s had increased proliferation rates (Ki67+) in ST2-ILC2s but not ST2+ ILC2s (**Figure 5J**). We next examined if gut LT-induced population-wise reduction of ST2 in lung ILC2s affected allergic lung responses. To directly test that, we used intranasal sensitization of LT-exposed mice with IL-33 (**Figure 6A**), which mimics lung type 2 allergic responses and is heavily dependent on ILC2 activation^15^. LT-exposed mice were significantly protected from intranasal IL-33 challenge: they displayed reduced BAL immune cell numbers (**Supplementary Figure 7**), diminished lung infiltration of total immune cells and eosinophils (**Figure 6B**) and developed significantly less tissue inflammation (**Figures 6C-D**). Altogether, these results suggest that gut ETEC/LT exposure induces the accumulation of lung ILC2s with reduced ability to respond to IL-33 and induce lung type 2 allergic inflammation.

**Figure 6:**
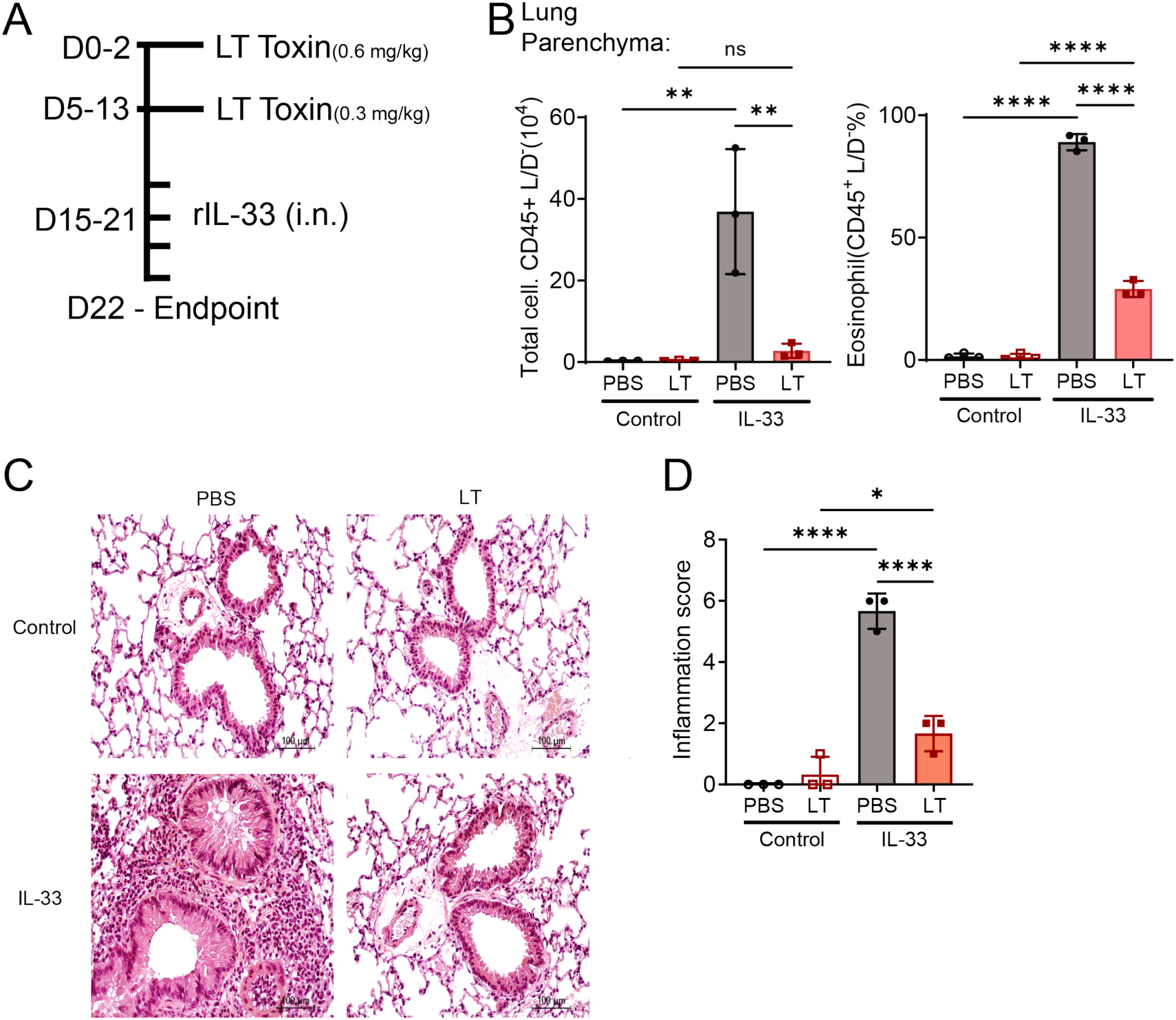
Gut LT exposure protects from IL-33 induced lung type 2 inflammation. (A) Representative scheme of the experimental timeline for oral gavage with LT and subsequent intranasal IL-33 inoculation. (B) Average numbers of live leukocytes and eosinophils in the lung parenchyma. (C) Representative lung sections stained with HE and PAS. (D) Average lung inflammation score values based on histological analyses. Data are shown as means ± SD (n = 3-5 mice) from 2 independent experiments. One-way ANOVA with Tukey’s multiple comparison test. ****P < 0.0001, ***P < 0.001, **P < 0.01, *P < 0.05.

### ETEC/LT-mediated release of gut IL-33 promotes protection from allergic lung inflammation

In contrast to reduced responses to IL-33 in the lung, ETEC/LT exposure led to increased IL-33 production and expression of ST2 by ILC2s in the gut (**Figures 5B-C**). Gut IL-33 release and subsequent activation of ILC2s have been linked to tissue repair and immune regulation^11,12^, which led us to question whether LT-induced type 2 response in the gut could paradoxically inhibit type 2 allergic inflammation in the lung. First, we infected IL-33KO mice with ETEC, followed by exposure to OVA+Pap (**Supplementary Figure 8A**). In comparison to WT mice, previously ETEC-infected IL-33KO mice displayed increased numbers of total immune cells and eosinophils in the BAL, and no reduction of these cells induced by ETEC (**Supplementary Figures 8B**), suggesting a lack of ETEC effect in the absence of IL-33. In addition, lung tissue accumulation of eosinophils, CD4+ Th2 cells and ILC2s were not altered by ETEC infection in IL-33KO mice (**Supplementary Figures C**). Because global deletion of IL-33 could lead to confounding effects in these experiments, we next used gut epithelial cell-specific IL-33KO mice (Villin-Cre IL33 fl/fl), exposing these mice with purified LT protein (**Figure 7A**). Gut epithelial-specific ablation of IL-33 had no effects on the development of OVA+Pap allergic lung inflammation, with similar numbers of BAL total cell and eosinophil accumulation (**Figures 7B-C**). Strikingly, pre-exposure to LT did not have any lung protective effects on OVA+Pap-exposed Villin-Cre IL33 fl/fl mice, and these mice had significantly higher levels of allergic lung inflammation if compared to LT-exposed WT mice (**Figures 7B-C**). Overall, these data suggest that gut epithelial cell-derived IL-33 accumulation following exposure to ETEC/LT is required for the acquisition of protection from allergic lung inflammation.

**Figure 7:**
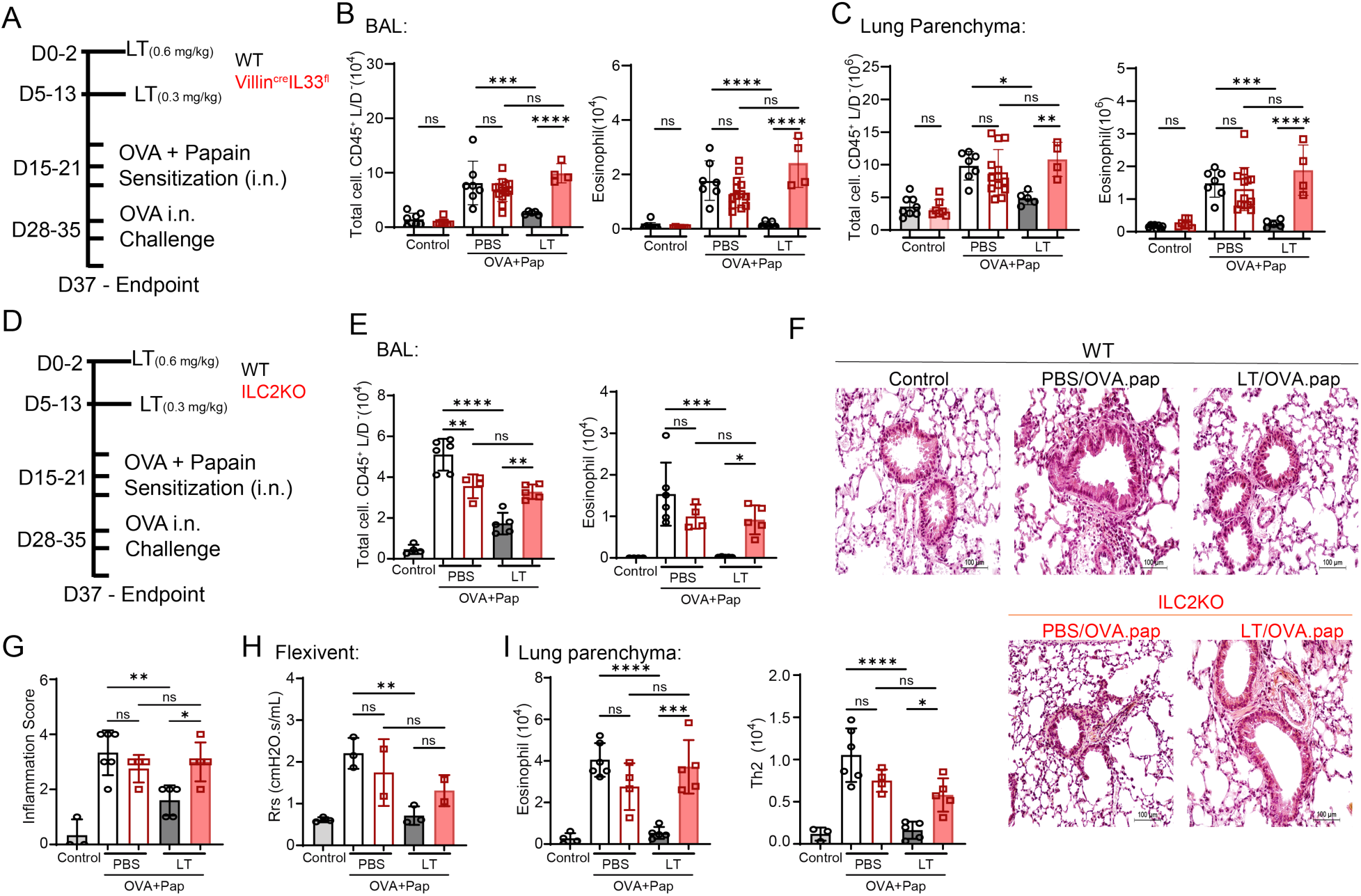
ETEC/LT-mediated protection from lung allergy is dependent on IL-33 and ILC2s. (A) Representative scheme of the development of lung allergy with OVA+Pap after oral gavage ETEC infection of WT and Villin-Cre IL33 fl/fl mice. (B) Average numbers of total live leukocytes, frequency and numbers of eosinophils in the BAL (C) Average numbers of total live leukocytes, frequency and numbers of eosinophils in the Lung parenchyma. (D) Representative scheme of the development of lung allergy with OVA+Pap after oral gavage ETEC infection of WT and ILC2KO mice. (E) BAL average numbers of total leukocytes and eosinophils in ETEC or PBS-exposed WT versus ILC2KO mice. (F) Representative lung sections of the indicated experimental groups, stained with H&E and PAS. (G) Average lung inflammation score values based on histological analyses. (H) Average values for lung respiratory system resistance (measured by oscillometry) in WT versus ILC2KO mice. (I) Average numbers of lung eosinophils and CD4+ Th2 cells in WT versus ILC2KO mice. Data are shown as means ± SD (n = 2 - 13 mice) from 2 independent experiments. One-way ANOVA with Tukey’s multiple comparison test. ****P < 0.0001, ***P < 0.001, **P < 0.01, *P < 0.05.

Because gut ILC2s accumulate and upregulate ST2 following ETEC/LT exposure, we tested if gut ILC2s would participate in the distal protection effect induced by gut IL-33. We used LT exposure followed by OVA+Pap allergy induction in IL7r-Cre Rorα fl/fl (ILC2KO) mice (**Figure 7D**). In comparison to WT mice, LT-exposed ILC2KO mice had increased BAL accumulation of eosinophils, and no reduction of BAL cellularity induced by LT (**Figures 7E, Supplementary Figure 8D-E**). Moreover, LT-induced protection from lung inflammation did not occur in ILC2KO mice (**Figures 7F-G**), which was reflected in their lack of improve in respiratory capacity in comparison to LT-exposed WT mice (**Figure 7H**). Lack of protection from lung inflammation was accompanied by no LT effects on the lung accumulation of total immune cells, eosinophils and CD4+ Th2 cells in ILC2KO mice, in comparison to WT mice (**Figures 7I**); predictably, no ILC2s were present in ILC2KO mice (**Figure Supplementary Figure 8F**). Type 2 inflammation induced by OVA+Pap was reduced but not absent in ILC2KO mice (**Figures 7E-I**), possibly due to the function of ST2^+^ CD4^+^ Th2 cells. Indeed, gut LT exposure leads to decreased ST2^+^ CD4^+^ Th2 cells in the lungs following OVA+Pap; this reduction, however, did not occur in ILC2KO mice (**Supplementary Figure 8G**). Overall, these results indicate that protection from lung allergy induced by ETEC/LT gut exposure requires the presence of ILC2s.

## Discussion

In this report, we define a non-canonical mechanism of negative regulation of lung type 2 allergic inflammation, induced by gut exposure to enterotoxigenic *E. coli* or its toxin LT and subsequent gut IL-33 release. Allergen-induced lung type 2 inflammation can be modulated by lung-resident immune cells and processes^46,56^. In contrast, much less is known about systemic or distal effects regulating these responses. Our work unveils a mechanism for gut-localized immune responses to reduce lung type 2 inflammation, constituting a distal negative feedback loop axis.

Paradoxically, *E. coli* LT induction of IL-33 release in the gut was accompanied by a reduction in the IL-33 sensitivity by lung ILC2 cells. The mechanisms by which this remote modulation occurs remain elusive. One possibility could be the recruitment of ST2^low^ ILC2s from other sites such as the bone marrow or the gut itself, like what has been recently reported to occur in other disease contexts^57–59^. Our CFSE tracing experiments, however, make this hypothesis unlikely, at least when it comes to rapid ILC2 mobilization. More precise fate mapping models will help with a definitive answer to this question. Another hypothetical mechanism could be the production and release of soluble factors by gut IL-33-responding cells. The ability of ST2-expressing gut cells to respond to local damage and produce cytokines or epidermal growth factor receptor such as IL-5, IL-10, IL-13 and amphiregulin as well as their potential effect in systemic immunity has been documented^6,60,61^. Whether such effects can be regulated by LT toxins remains to be determined and would likely mean gut exposure to these toxins regulate immune responses in multiple organs, beyond the effects observed in the lungs.

Alternatively, it is also possible that gut IL-33 is driving remote changes in lung immune responses through the action of enteric nervous system cells. Indeed, several ST2-expressing gut cells typically co-localize with sensory and parasympathetic enteric neurons, such as ILC2s^62–64^. Future experiments targeting these neurons specifically will be necessary to test whether *E. coli* toxin exposure and IL-33 release drives systemic immune changes through a gut-nervous system axis. If this is true, it will also be important to determine how gut IL-33-mediated neuronal signals can modulate lung ILC2 responses. Recent reports showed that lung ILC2 cells can respond directly, for example, to dopamine released locally, with an anti-allergenic effect^65^. In future studies, determining whether gut IL-33 release negatively regulates lung ILC2 allergy responses through dopaminergic signaling may help define how hosts are protected from lung allergy following *E. coli* infection and LT exposure.

Regardless of the mechanism, the observed dependency on IL-33 release strongly indicates that gut ST2-expressing cells play a fundamental role in the regulation of lung type 2 allergy. Multiple immune and non-immune cells in the small intestine can express ST2, such as Tuft cells, mast cells, Th2 CD4^+^ T cells, regulatory T cells or ILC2s^66–68^. Despite not excluding the possibility of multiple cell types performing this function, our current work indicates that gut ILC2s, which upregulate ST2 following *E. coli* or LT, may play a relevant role in the downregulation of lung type 2 inflammation. Future research using systems to specifically ablate gut ILC2s will be necessary to corroborate this working hypothesis, as well as to further define how these cells can modulate lung type 2 inflammation.

Overall, our study suggests that prior exposure to pathogenic bacterial infections such as *E. coli* has an unanticipated protective effect against allergic lung inflammation, through an atypical gut IL-33-mediated effect. These data help provide a mechanistic explanation for an underappreciated angle on the so-called “hygiene hypothesis”, which is the inverse correlation between the incidence of diarrhea-inducing infectious diseases and lung allergy^8,20,32^. More importantly, it opens the possibility of using oral exposure to IL-33 inducing molecules as a therapeutic avenue to control asthma or other allergic lung diseases.

## Abbreviations

CEUA: *Comissão de Ética no Uso de Animais* (Ethics Committee for Animal Use, USP)
DTT: Dithiothreitol
EDTA: Ethylenediaminetetraacetic acid
ETEC: Enterotoxigenic *Escherichia coli*
FACS: Fluorescence-activated cell sorting
GM1: Monosialotetrahexosylganglioside
gMFI: Geometric mean fluorescence intensity
H&E: Hematoxylin and eosin staining
IACUC: Institutional Animal Care and Use Committee
IL-33KO: Interleukin-33 knockout
ILC2KO: Group 2 innate lymphoid cell–deficient mice (Il7r-Cre × Rorα flox/flox)
KO: Knockout
LD: Live/Dead dye
LT: Heat-labile toxin (enterotoxin from *E. coli*)
LTB: Heat-labile toxin subunit B
LTK63: Non-toxic mutant of heat-labile toxin
LT-KO ETEC: Plasmid-cured non-toxinogenic ETEC (LT^−^)
Pap: Papain
PAS: Periodic acid–Schiff staining
PMA: Phorbol myristate acetate
SCIREQ: Scientific Respiratory Equipment Inc.
SD: Standard deviation
siLP: small intestine lamina propria
WT: Wild type

## Notes

Funding Statement: This study was supported by São Paulo Research Foundation (FAPESP) grants: 2021/06881-5, 2021/15185-2, 2019/13916-0 and 2015/25364-0; National Council for Scientific and Technological Development (CNPq) grant 315712/2023-6; and the National Institutes of Health/National Institute of Allergy and Infectious Diseases (NIH/NIAID grant AI170649).

Conflict of interest statement: The authors have declared that no conflict of interest exists

### Competing Interest Statement

The authors have declared no competing interest.

