## Supplementary Material for "*Escherichia coli*-induced gut IL-33 release inhibits lung type 2 allergic responses"

| Antibody | Company | Catalog | Antibody | Company | Catalog |
|---|---|---|---|---|---|
| B220 | Ebioscience | 47-0452-82 | IL-5 | Biolegend | TRFK5 |
| CCR2 | R&D | FAB5538P-100 | IL-13 | Ebisocience | Ebio13a |
| CD4 | Biolegend | 100547 | IL-17 | Ebioscience | Ebio17B7 |
| CD8b | Biolegend | YTS156.7.7 | IL-22 | Biolegend | Poly5164 |
| CD11b | Biolegend | 101257 | Live and Dead | Invitrogen | N/A |
| CD11c | Biolegend | 117336 | Ly6C | Biolegend | HK1.4 |
| CD44 | Biolegend | IM7 | Ly6G | Biolegend | 127626 |
| CD45 | Biolegend | 103151 | Ki67 | Biolegend | 652426 |
| CD64 | Biolegend | X54-5/7.1 | MHC II | Biolegend | M5/114.15.2 |
| CD103 | Ebioscience | 46-1031-82 | NK1.1 | Biolegend | 108736 |
| CD206 | Biolegend | C068C2 | RORvt | Ebioscience | 46-6981-82 |
| FC Block (anti-CD16/32) | Bioxcell | 2.4G2 | SiglecF | BD Biosciences | 562757 |
| FOXP3 | Ebioscience | 12-5773-82 | Tbet | Ebioscience | EBIO4B10 |
| GATA3 | BD Biosciences | 560077 | TCRb | Ebioscience | 47-5961-82 |
| IL-4 | Biolegend | 11B11 | Thy1.2 | Biolegend | 105331 |
| ST2 | Biolegend | DIH4 | | | |

**Supplementary Table 1:** List of antibodies, company and reference catalog.

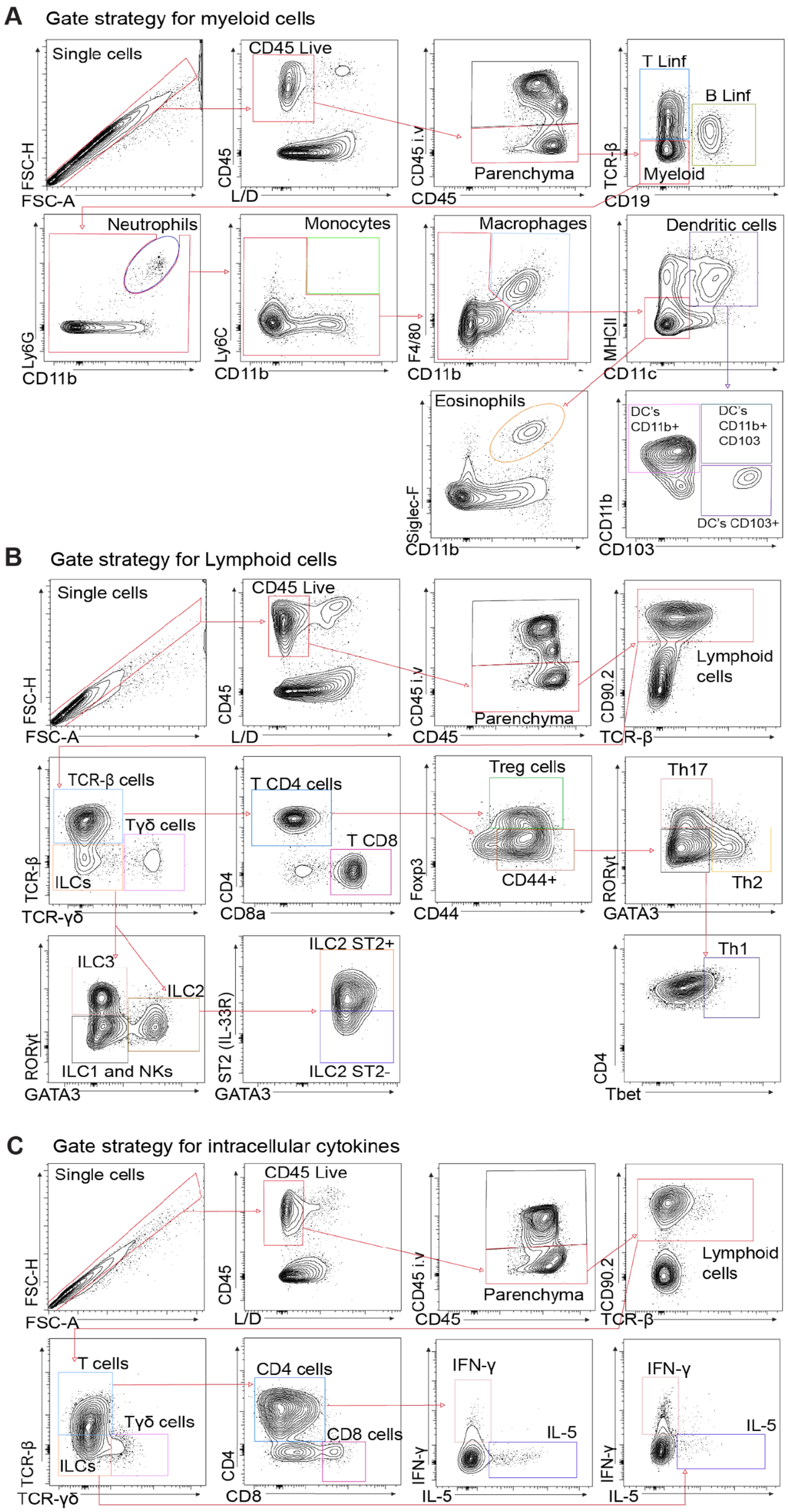
A
Gate strategy for myeloid cells
Single cells
FSC-H
FSC-A
CD45 Live
CD45
L/D
CD45 i.v
CD45
Parenchyma
T Linf
B Linf
TCR-β
Myeloid
CD19
Neutrophils
Ly6G
CD11b
Monocytes
Ly6C
CD11b
Macrophages
F4/80
CD11b
Dendritic cells
MHCII
CD11c
Eosinophils
Siglec-F
CD11b
DC's CD11b+
DC's CD11b+ CD103
DC's CD103+
CD11b
CD103
B
Gate strategy for Lymphoid cells
Single cells
FSC-H
FSC-A
CD45 Live
CD45
L/D
CD45 i.v
CD45
Parenchyma
Lymphoid cells
CD90.2
TCR-β
TCR-β cells
Tγδ cells
ILCs
TCR-β
TCR-γδ
T CD4 cells
T CD8
CD4
CD8a
Treg cells
CD44+
Foxp3
CD44
Th17
Th2
RORγt
GATA3
ILC3
ILC2
ILC1 and NKs
RORγt
GATA3
ILC2 ST2+
ILC2 ST2-
ST2 (IL-33R)
GATA3
Th1
CD4
Tbet
C
Gate strategy for intracellular cytokines
Single cells
FSC-H
FSC-A
CD45 Live
CD45
L/D
CD45 i.v
CD45
Parenchyma
Lymphoid cells
CD90.2
TCR-β
T cells
Tγδ cells
ILCs
TCR-β
TCR-γδ
CD4 cells
CD8 cells
CD4
CD8
IFN-γ
IL-5
IFN-γ
IL-5
IFN-γ
IL-5
IFN-γ
IL-5

2

**Supplementary Figure 1. Gating strategies.** Flow cytometry gating strategies for detection in lung and small intestine of: (A) myeloid cells – neutrophils, monocytes, macrophages, dendritic cells and eosinophils; (B) lymphoid cells – $CD8^+$ T cells, $CD4^+$ T cell subsets (Th1, Th2, Th17, Treg cells), γδ T cells, ILC1/NK cells, ILC2s and ILC3s; (C) staining for intracellular cytokines – IFN-γ and IL-5.

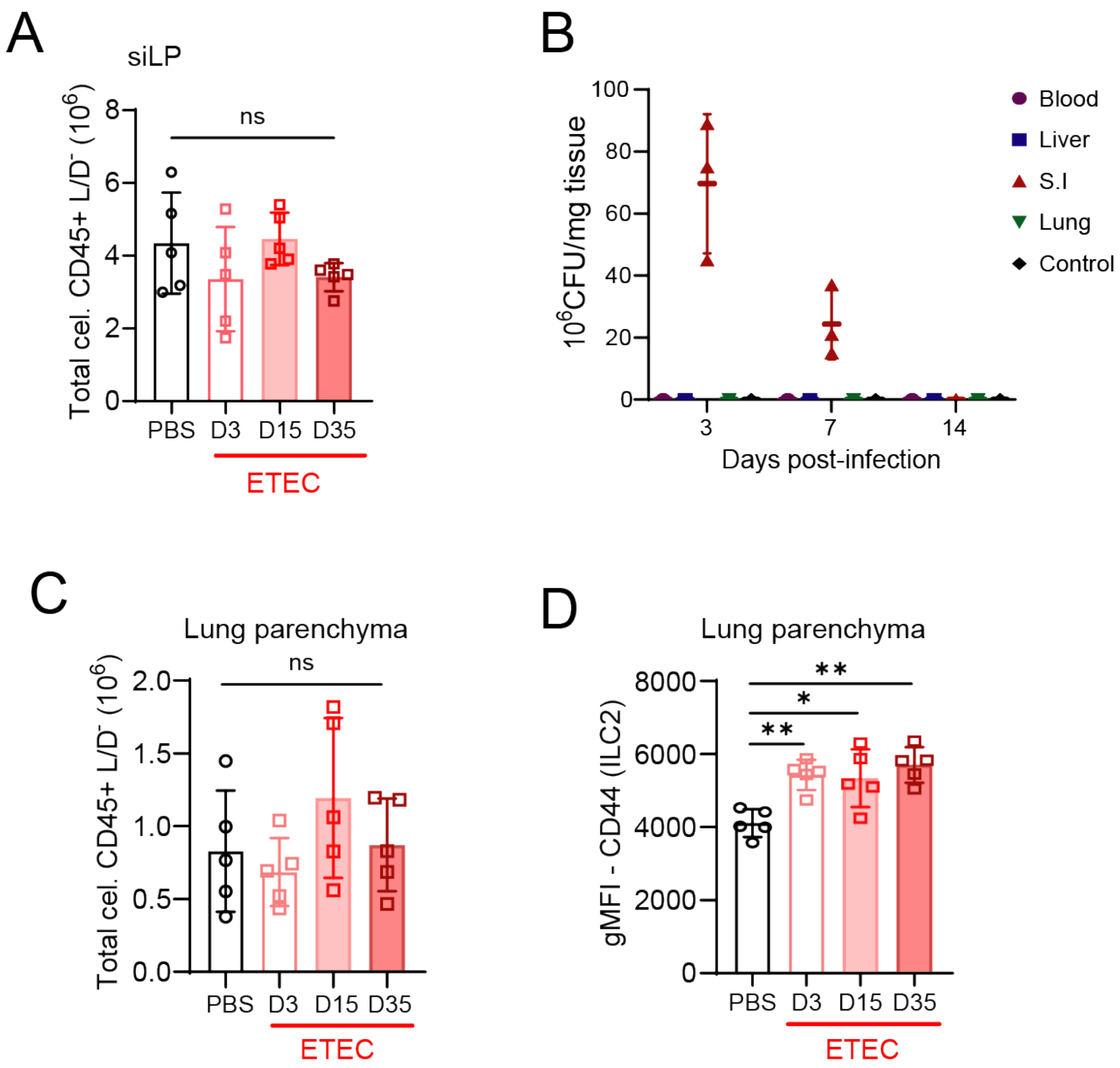


**Supplementary Figure 2. Effects of ETEC infection in the small intestine and lungs.** WT mice were orally inoculated with $10^9$ ETEC bacteria. Multiple organs were harvested and analyzed at days 3, 15 and 35 p.i.. (A) Average number of $CD45^+$ leukocytes in the small intestine lamina propria (siLP). (B) Colony forming units (CFU) of ETEC in the indicated organs. (C) Average number of $CD45^+$ leukocytes in the lung. (D) Average gMFI values for CD44 expression by ILC2s in the lung parenchyma. Data are shown as means ± SD (n = 3-6 mice) from 2 independent experiments. One-way ANOVA with Tukey's multiple comparison test. ns – not significant, $^{**}P < 0.01$, $^{*}P < 0.05$.

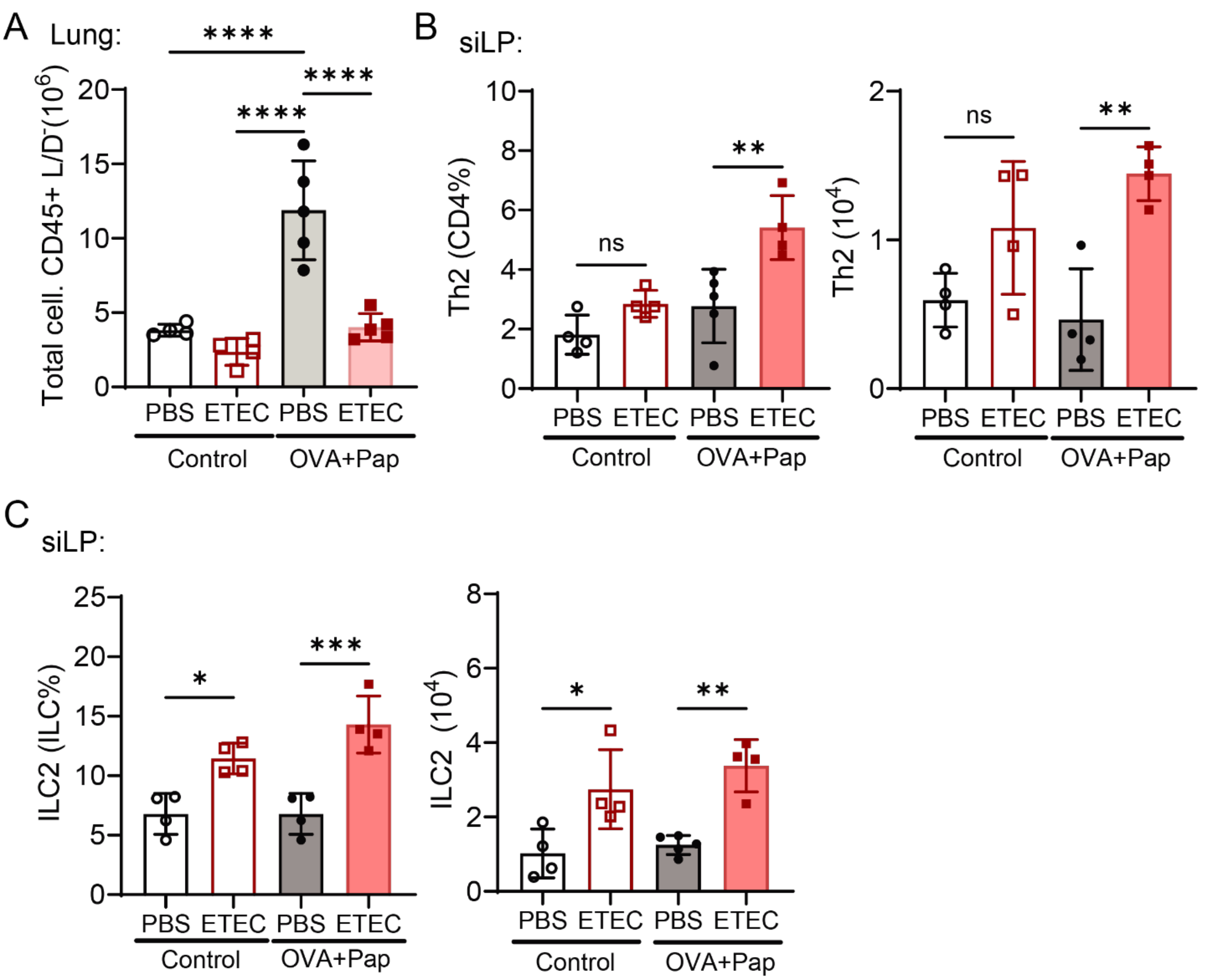


**Supplementary Figure 3. Effects of ETEC infection and allergy induction in the small intestine and lungs.** WT mice were pre-exposed to ETEC, then sensitized and challenged intranasally with OVA+Papain. (A) Average numbers of $CD45^+$ leukocytes in the lung. (B) Average percentages (left) and numbers (right) of Th2 cells in the small intestine lamina propria (siLP). (C) Average percentages (left) and numbers (right) of ILC2s in the siLP. Data are shown as means ± SD (n = 4-5 mice) from 2 independent experiments. One-way ANOVA with Tukey's multiple comparison test. ns – not significant, ****P<0.0001, ***P<0.001, **P < 0.01, *P < 0.05.

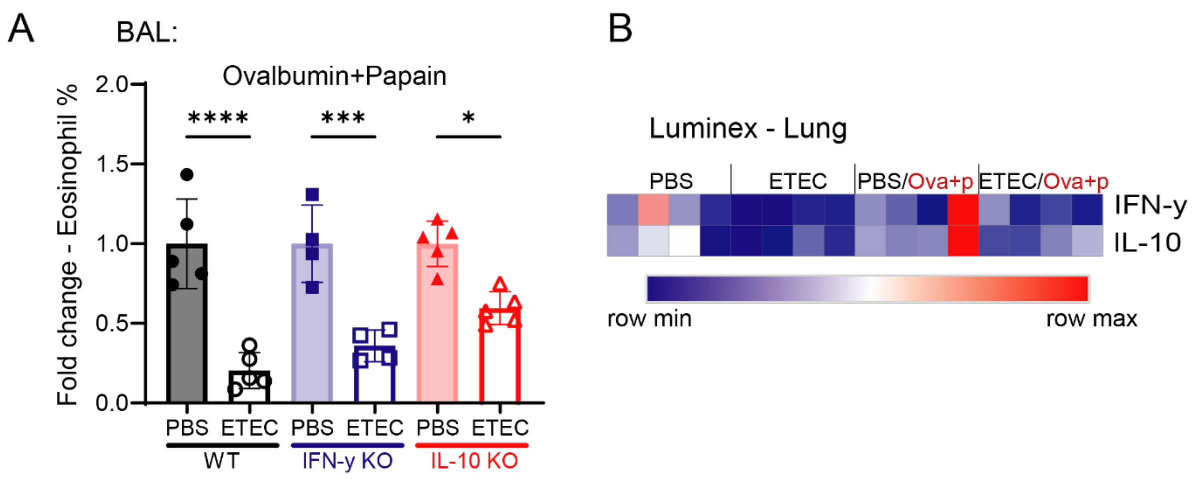


**Supplementary Figure 4. ETEC-mediated protection from lung allergy is independent of IFN-γ or IL-10.** WT, IFN-γ KO or IL-10 KO mice were pre-exposed to ETEC, then sensitized and challenged intranasally with OVA+Pap. (A) Average fold changes in eosinophil percentages between PBS-treated (control) and ETEC-treated mice in the lung bronchoalveolar lavage (BAL). (B) Heatmap of cytokine levels (IFN-γ, IL-10) measured in lung tissues of WT mice using Luminex assay. Data are shown as means ± SD (n = 4-5 mice) from 2 independent experiments. One-way ANOVA with Tukey's multiple comparison test. ****P < 0.0001, ***P < 0.001, *P < 0.05.

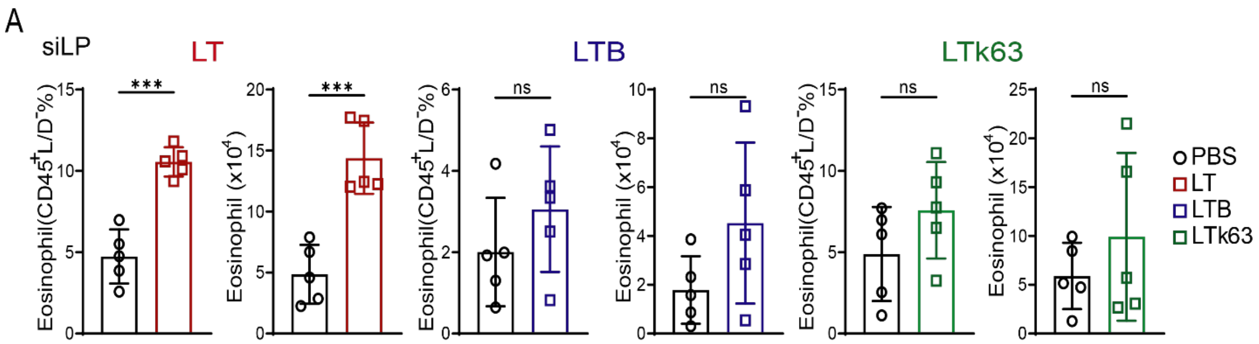


**Supplementary Figure 5. Eosinophilic response in the siLP exposed to different heat-labile toxin subtypes.** WT mice were orally exposed to LT, LTB or LTk63. Average frequencies and numbers of eosinophils in the siLP exposed to LT, LTB or LTk63. Data are shown as means ± SD (n = 5 mice) from 3 independent experiments. One-way ANOVA with Tukey's multiple comparison test. ***$P < 0.001$.

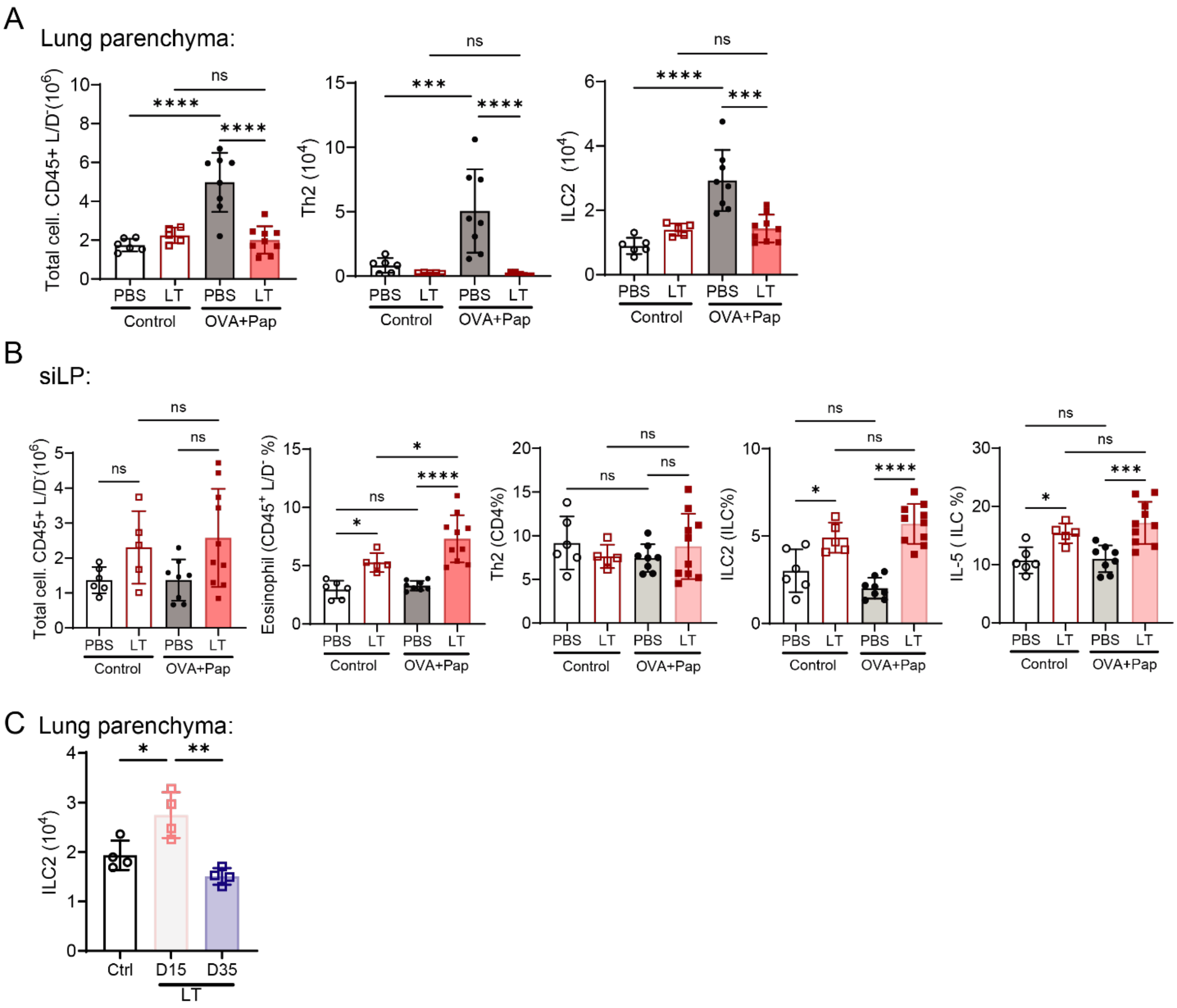


**Supplementary Figure 6. Gastrointestinal exposure to LT toxin is sufficient to promote protection from type 2 lung inflammation through the development of a type 2 response in the intestine.** WT mice were orally exposed to LT, then sensitized and challenged intranasally with OVA+Pap. (A) Average numbers of total leukocytes (CD45+ L/D-) and average numbers of Th2 and ILC2s in the lung parenchyma. (B) Average numbers of total leukocytes (CD45+ L/D-) and average frequencies of eosinophils, Th2, ILC2s and IL-5-producing ILCs in the siLP. (C) Kinetic of ILC2 recruitment to the lung of WT mice following oral exposure to LT. Data are shown as means ± SD (n = 4-9 mice) from 3 independent experiments. One-way ANOVA with Tukey's multiple comparison test. ****P < 0.0001, ***P < 0.001, *P < 0.05.

BAL:

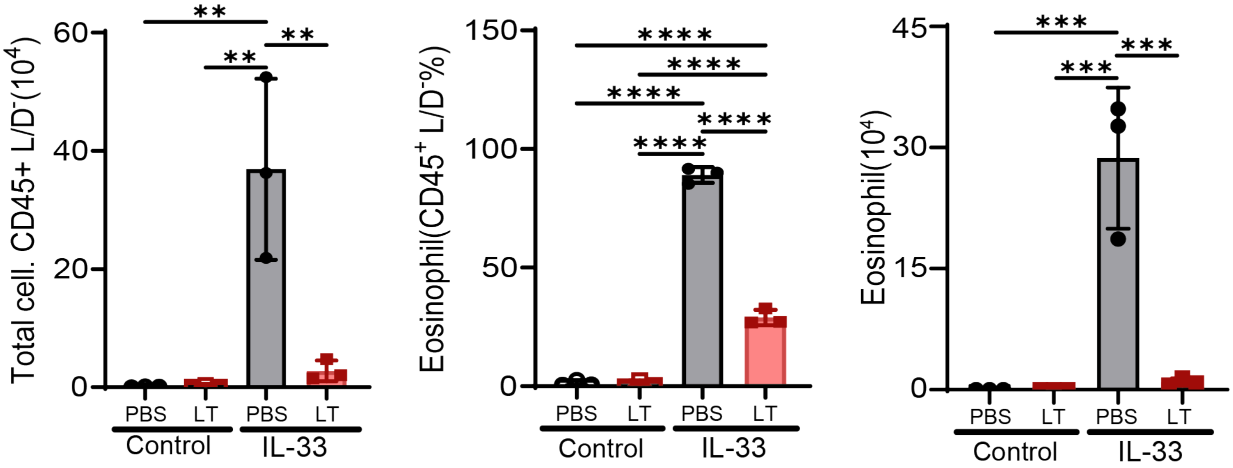


**Supplementary Figure 7. Gastrointestinal LT exposure protects from IL-33-induced eosinophilic response in the BAL.** WT mice were orally exposed to LT, then sensitized intranasally with recombinant IL-33. (A) Average numbers of total leukocytes and frequency and numbers of eosinophils in the BAL. Data are shown as means ± SD 3-5 mice) from 2 independent experiments. One-way ANOVA with Tukey's multiple comparison test. ****P < 0.0001, ***P < 0.001, **P < 0.01.

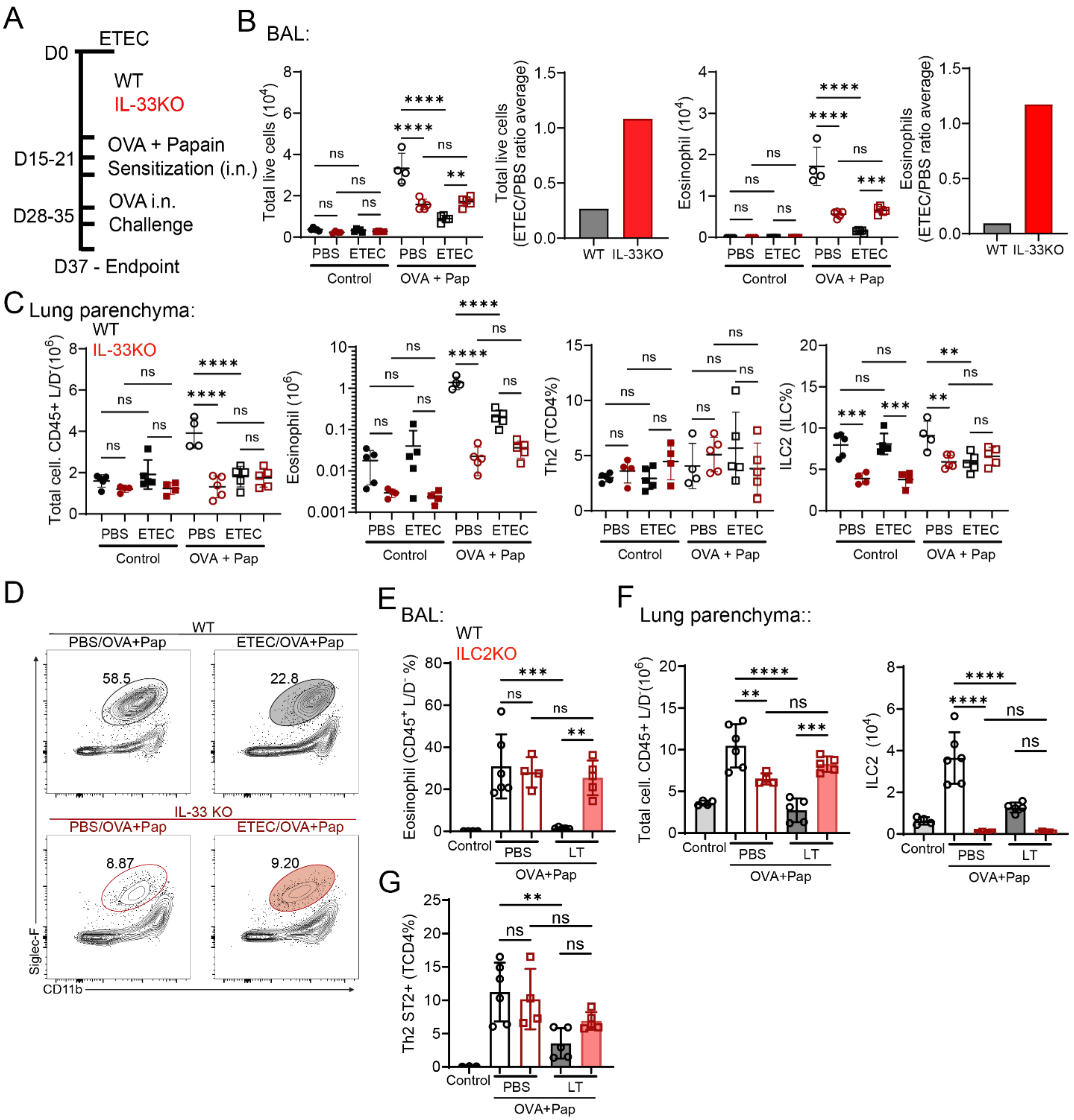


**Supplementary Figure 8. ETEC/LT-mediated protection from type 2 inflammation in the lung is dependent on IL-33 and ILC2s.** WT and IL-33KO mice were pre-exposed to ETEC, then sensitized and challenged intranasally with OVA+Pap. (A) Representative scheme of the development of lung allergy with OVA+Pap after ETEC infection of WT and IL-33KO mice. (B) In left-to-right sequence: BAL average numbers of total leukocytes; the ETEC/PBS ratios for total leukocytes in WT versus IL-33KO mice; BAL average numbers of eosinophils; and the ETEC/PBS ratios for eosinophils in WT versus IL-33KO mice. (C) Average numbers of total leukocytes (CD45+ L/D-) and average frequency of total leukocytes, eosinophils, Th2 and ILC2s in the lung of IL-33KO and WT mice. (D) Representative flow cytometry plots showing WT and IL-33KO's lung parenchyma eosinophils. Percentages are based on the gating strategy from Supplementary Figure 1. (E). Average numbers of total leukocytes (CD45+ L/D-) in the BAL of ILC2KO and WT mice. (F) Average numbers of total leukocytes (CD45+ L/D-) and average numbers of ILC2s in the lung parenchyma of ILC2KO and WT mice. (G) Average frequency of Th2 ST2+ in the lung of ILC2KO and WT mice. Data are shown as means ± SD (n = 2 - 5 mice) from 2 independent experiments. One-way ANOVA with Tukey's multiple comparison test. ****P < 0.0001, ***P < 0.001, **P < 0.01.
